# The evolution of bacterial shape complexity by a curvature-inducing module

**DOI:** 10.1101/2020.02.20.954503

**Authors:** Nicholas R. Martin, Edith Blackman, Benjamin P. Bratton, Thomas M. Bartlett, Zemer Gitai

## Abstract

Bacteria can achieve a staggering diversity of cell shapes that promote critical functions like growth, motility, and virulence^1-4^. Previous studies suggested that bacteria establish complex shapes by co-opting the core machineries essential for elongation and division^5,6^. In contrast, we discovered a two-protein module, CrvAB, that can curve bacteria autonomously of the major elongation and division machinery by forming a dynamic, asymmetrically-localized structure in the periplasm. CrvAB is essential for curvature in its native species, *Vibrio cholerae*, and is sufficient to curve multiple heterologous species spanning 2.5 billion years of evolution. Thus, modular shape determinants can promote the evolution of morphological complexity independently of existing cell shape regulation.

## Main Text

Cells come in a variety of shapes that contribute to fitness in diverse environments ^1,2,7,8^. In bacteria, cell curvature can influence motility and host colonization ^1,3,4^. While bacteria can achieve a huge diversity of shapes, all of these shapes are defined using a rigid cell wall made from a meshwork of peptide-crosslinked polysaccharide strands, termed peptidoglycan (PG)^9^. The mechanisms by which bacteria achieve complex shapes remain largely mysterious, but in all cases studied to date they involve species-specific co-opting of the machinery required for making simple shapes ^5,6^. For example, MreB, a cytoskeletal organizer required for the formation of simple rod shapes in most bacterial species ^10,11^, is required for the elaboration of both curvature and stalk formation in *C. crescentus* ^5,6^. In addition to MreB, the other widely-conserved cytoskeletal organizer is FtsZ ^12,13^. Each of these cytoskeletal elements localizes asymmetrically to direct the activity of an associated set of PG synthesis enzymes. Here we describe a cell curvature module that can function in multiple species independently of both the MreB- and FtsZ-associated “core” shape machineries, demonstrating that the evolution of bacterial shape is more plastic than previously appreciated.

### Two genes, *crvA* and *crvB*, are sufficient to induce cell shape complexity

To understand the origin of a complex curved-rod shape, we focused on the human pathogen *Vibrio cholerae*. The only known factor required for cell curvature in *V. cholerae*, CrvA, forms a protein filament that localizes to the inner face of the cell to induce cell curvature ^14^. The sequence downstream of the *crvA* gene encodes a larger gene with homology to *crvA*, which we named *crvB* (Fig. S1A, B). To determine whether this second gene is also a curvature determinant, we deleted it and found that the resulting cells were straight (Fig. 1A). Complementation tests showed that *crvA* complemented *ΔcrvA* and *crvB* complemented *ΔcrvB*, but *crvA* and *crvB* could not cross-complement (Fig. S1C, D, E). We searched for CrvB homologs in the genomes of other *Vibrio* species and found that almost every *Vibrio* with a CrvA homolog also has a CrvB homolog (Fig. 1B). The singular exception, *Vibrio proteolyticus*, contains only a CrvA homolog and is reported to be a straight rod ^15^. Furthermore, many species have neither a CrvA nor a CrvB homolog and are reported to be straight. Thus, not only are CrvA and CrvB both required for cell curvature in *V. cholerae*, but their evolutionary co-occurrence correlates with cell curvature throughout the *Vibrio* genus.

**Fig. 1.**
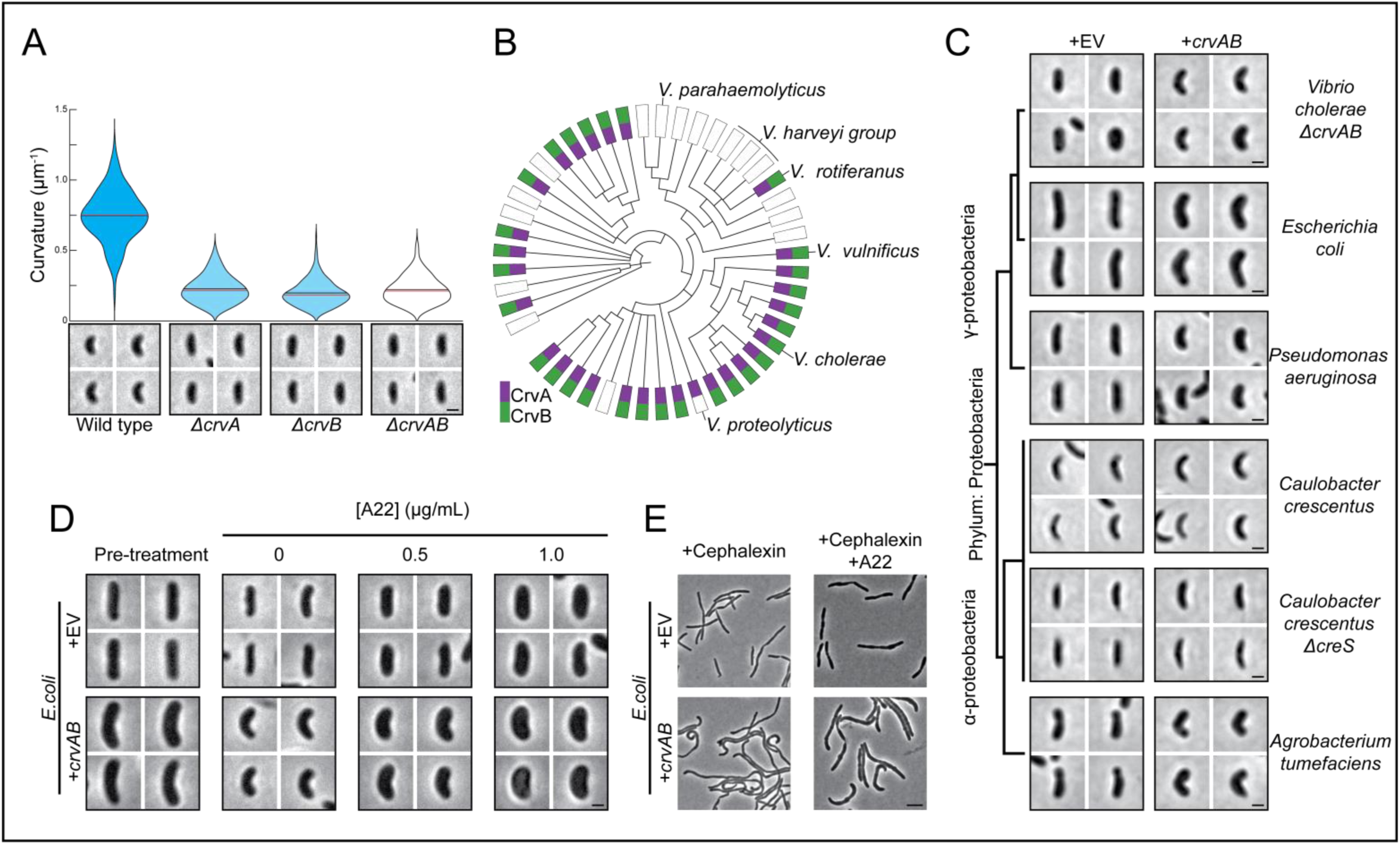
CrvA and CrvB curve bacterial cells. (A) Single-cell curvature measurements using Morphometrics ^10^ from indicated *V. cholerae* strains. Bars represent mean (black) and median (red). (B) Presence of CrvA and CrvB homologs across *Vibrio* species. (C-E) Heterologous expression of *crvA* and *crvB* (+*crvAB*) or empty vector (+EV). (C) Brackets indicate phylogenetic relationships between species. (D) *E. coli* before (Pre-treatment) and after A22 treatment (E) *E. coli* after treatment with cephalexin and A22. (A,C,D) Images represent 95^th^ percentile of curvature in respective populations. Scale bars are 1μm. (B) Clade structure drawn according to ^26^. (C) Median curvature of +crvAB is significantly higher than +EV in each species (p≤3.74×10^−5^). (D) Median curvature of +crvAB is significantly higher than +EV and Pre-treatment at each A22 concentration (p≤2.02×10^−9^). Wilcoxon rank sum tests used for all statistics; n=300 (See FigS2 for quantified populations). Scale bars represent 1μm in (A,C,D) and 5μm in (E).

To determine whether CrvA and CrvB are sufficient to induce curvature, we introduced them into the straight-rod species *Escherichia coli* and *Pseudomonas aeruginosa*. Despite the fact that *crvA/crvB* homologs are restricted to the family *Vibrionaceae*, heterologous expression was sufficient to transform both *E. coli and P. aeruginosa* into curved rods. Thus, *crvA* and *crvB* represent the complete set of *V. cholerae-*specific genes required for curvature (Fig. 1C, Fig. S2). We next sought to determine if CrvA and CrvB are sufficient to induce curvature in a species that is not normally a straight rod. *V. cholerae, E. coli*, and *P. aeruginosa* are γ-proteobacteria, while the curved-rod *Caulobacter crescentus* is an α-proteobacterium. *C. crescentus* lacks *crvA/crvB* homologs but requires crescentin (*creS*) for curvature such that a *ΔcreS* mutant is a straight rod. CreS does not bear substantial primary sequence homology to CrvA or CrvB ^14^. Nevertheless, heterologous CrvA and CrvB expression curved *ΔcreS* cells (Fig. 1C, Fig S2) and significantly enhanced cell curvature in wild-type *C. crescentus* (Fig. 1C, Fig. S2). Thus, CrvA and CrvB can function in *C. crescentus* independently of CreS.

### CrvA and CrvB form a module that functions autonomously of core shape machineries

While *V. cholerae, E. coli, P. aeruginosa*, and *C. crescentus* use the MreB-directed machinery, known as the elongasome ^5,10,16,17^, to insert new PG throughout the lateral surface of the cell, some bacteria form rods via a distinct growth pattern. For example, the α-proteobacterium *Agrobacterium tumefaciens* lacks many canonical elongasome members including MreB, and instead co-opts FtsZ-directed synthesis to elongate from the cell pole ^18,19^. Surprisingly, CrvA and CrvB expression induced cell curvature in *A. tumefaciens* despite its distinct spatial pattern of growth and the absence of MreB (Fig. 1C, Fig. S2). Because CrvA and CrvB are sufficient to induce curvature in multiple heterologous species, including both straight and curved species, as well as in species that use either MreB-driven lateral growth or FtsZ-mediated polar growth, we refer to them as the CrvAB cell shape module.

*V. cholerae* curvature is robust to inhibition of cell division ^14^ and the MreB-independent function of the CrvAB module in *A. tumefaciens* suggests that the elongasome is not required for induction of cell curvature. To further test the dependence of CrvAB on the core shape machineries, we inhibited the elongasome and the FtsZ-associated “divisome” machinery in *E. coli* expressing CrvAB. Treatment with the MreB-disruptor, A22, induced a dose-dependent widening of cells (Fig 1D, Fig. S3A) indicative of disrupted MreB function ^20^. Nonetheless, CrvAB expression increased cell curvature over the duration of A22 treatment at all concentrations tested (Fig. 1D, Fig. S3B). CrvAB expression also induced curvature during treatment with the divisome inhibitor, cephalexin, alone or in combination with A22 (Fig. 1E). The ability of the CrvAB module to function independently of both core machines demonstrates that the emergence of complex cell shape can be achieved by the horizontal transfer of only two genes that act autonomously of endogenous cell shape regulation.

### The evolution of the CrvAB cell shape module

CrvA and CrvB share sequence homology, but their inability to complement one another suggests that they may have distinct functions. We thus looked for differences in their amino acid sequences and found that CrvB contains a ∼250 amino acid C-terminal domain that is not present in CrvA. Deletion of either this <u>C</u>rv<u>B</u>-<u>s</u>pecific (CBS) domain or the <u>N</u>-terminal <u>d</u>omain (NTD) of CrvB abolished curvature (Fig S4A). Thus, the CBS domain is necessary, but not sufficient for CrvB function. Remarkably, when we fused the CBS domain to the C-terminus of CrvA, the resulting chimeric protein, CrvA_CBS_, induced curvature in a *ΔcrvAB* background (Fig. 2A, Fig S4A). In *V. cholerae*, cell curvature rapidly decreases when saturated cultures are diluted into fresh media and increases until the culture reaches stationary phase^14^. While *crvA*_*CBS*_ was sufficient to curve cells, these cells did not become as curved as wild type (Fig 2B). To determine if increasing the level of *crvA*_*CBS*_ could further increase curvature, we expressed a second copy of *crvA*_*CBS*_ at the *crvB* locus (2xCrvA_CBS_) and found that this strain reached nearly wild-type curvature (Fig. 2B). However, CrvA_CBS_ induced curvature slower than wild type even when two copies were present (Fig. 2B). Therefore, a CrvA/CrvB hybrid functions as a minimal CrvAB module but does so slower than the two separate CrvA and CrvB determinants found in wild type.

**Fig. 2.**
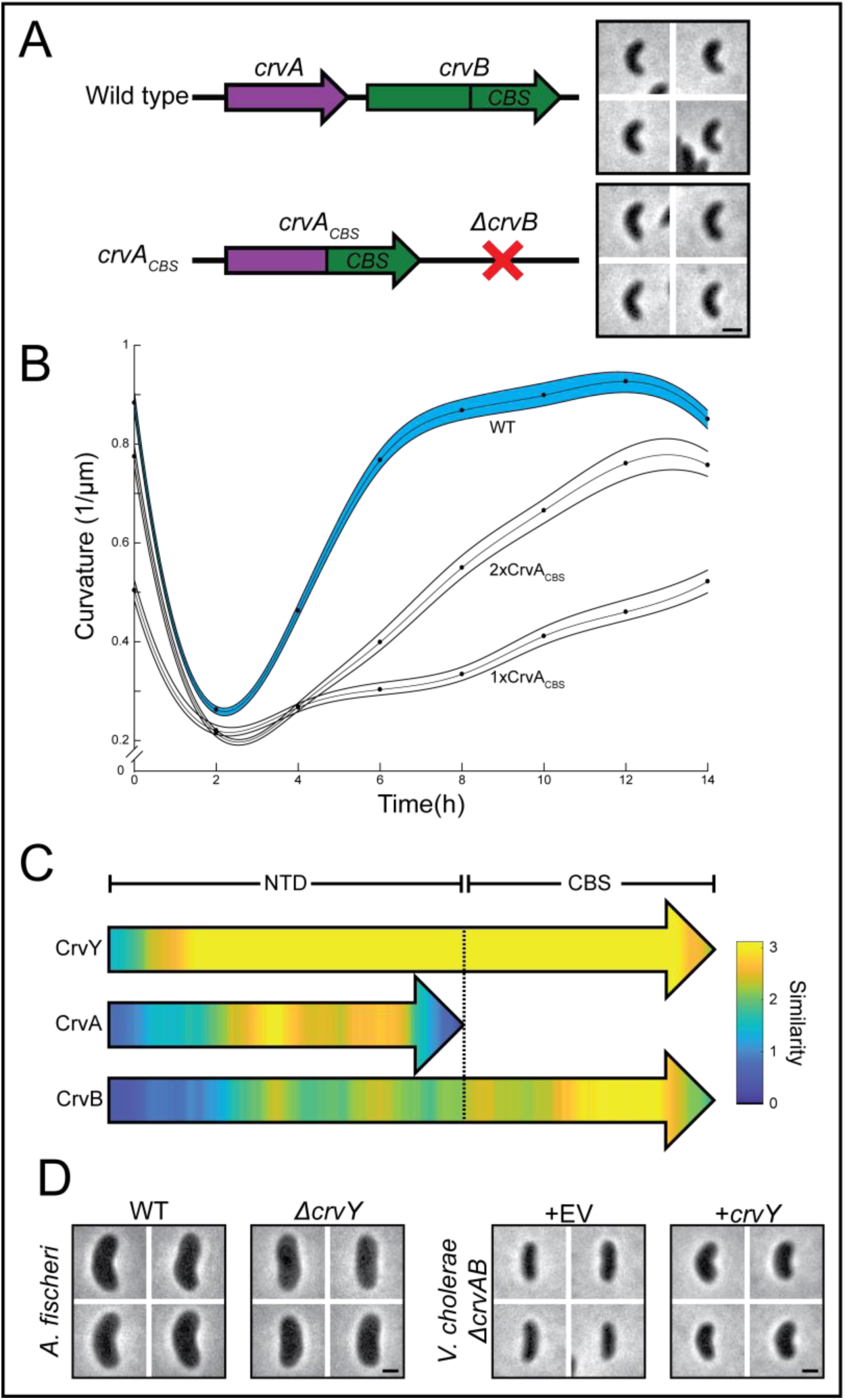
CrvA and CrvB are functionally and structurally specialized. (A) Schematic of the *crvAB* locus and representative images of wild type and cells expressing the *crvA*_*CBS*_ chimera. (B) Dynamics of mean curvature ± 95% confidence intervals (bootstrapped) in wild type (WT) and strains expressing one (1xCrvA_CBS_) or two (2xCrvA_CBS_) copies of *crvA*_*CBS*_. (C) Pairwise sequence alignment of all sequenced CrvA, CrvB, and CrvY homologs to CrvY. Sequence alignment and local similarity using the BLOSUM80 scoring matrix. (D) Deletion of *crvY* in *A. fischeri* and expression of *crvY*_*A. fischeri*_ in *V. cholerae ΔcrvAB*. (A,D) Images represent 95^th^ percentile of curvature in respective populations. Scale bars represent 1μm. See FigS4 for population measurements.

The division of functional domains among CrvA and CrvB suggested the possibility that the CrvAB module evolved from an ancestral protein resembling CrvA_CBS_. We performed a phylogenetic analysis of all sequenced CrvA/CrvB homologs and found a set of sequences that resemble a CrvA_CBS_ chimera in clades related to *Vibrios* such as *Allivibrios* (Figs. S4B-D). These proteins, which we refer to as CrvY, are distinct from both CrvA and CrvB, as they contain a CBS domain similar to that of CrvB but are more similar to CrvA in the rest of the protein (Fig 2C). Further phylogenetic analysis suggested that the common ancestor of species with CrvAB homologs had a single Crv protein that contained a CBS domain (Figs. S4B-D). To analyze the function of CrvY we focused on the curved species *Aliivibrio fischeri*. Deletion of *crvY* (*VF_A0834*) resulted in straight *A. fischeri* cells and heterologous expression of *crvY* in *V. cholerae* was sufficient to induce curvature in a *ΔcrvAB* mutant (Fig. 2D, Figs. S4E,F). The necessity of CrvY for curvature in *A. fischeri* and sufficiency for curvature in *V. cholerae* indicate that CrvY is a functional, extant form of a CrvA_CBS_ chimera. Our data thus suggest that the CrvAB module evolved by the duplication of a single ancestral gene followed by the loss of the CrvA CBS domain and divergence of the CrvB N-terminal domain.

### CrvAB forms a periplasmic filament that generates asymmetry and governs cell curvature dynamics

How do CrvAB induce an asymmetric cell shape without co-opting the core shape machinery? CrvA tagged with the fluorescent protein msfGFP (CrvA-GFP) is functional and assembles into a filamentous structure localized to the inner curvature of the cell ^14^. We found that a CrvB*-* msfGFP fusion was also functional and formed a filament at the inner curve (Fig. 3A, Fig. S5A). An msfGFP fusion to CrvA_CBS_ was functional and also localized in the same pattern (Figs. S4A, S5B). When we imaged mCherry-tagged CrvA (CrvA-mCherry) and CrvB-GFP simultaneously, both proteins localized to the same structure (Fig. 3A, Fig. S5A) and remained colocalized as cells grew (Fig. 3B, Movies S1-S2). Furthermore, CrvA-mCherry and CrvB-GFP colocalized upon heterologous expression in *E. coli*, indicating that CrvAB form an asymmetric structure in a species*-*independent manner (Fig. 3A, Fig. S5C). Thus, CrvA and CrvB are together sufficient to generate an asymmetric structure in *E. coli*, which could explain why they do not need MreB or FtsZ to break symmetry.

**Fig. 3.**
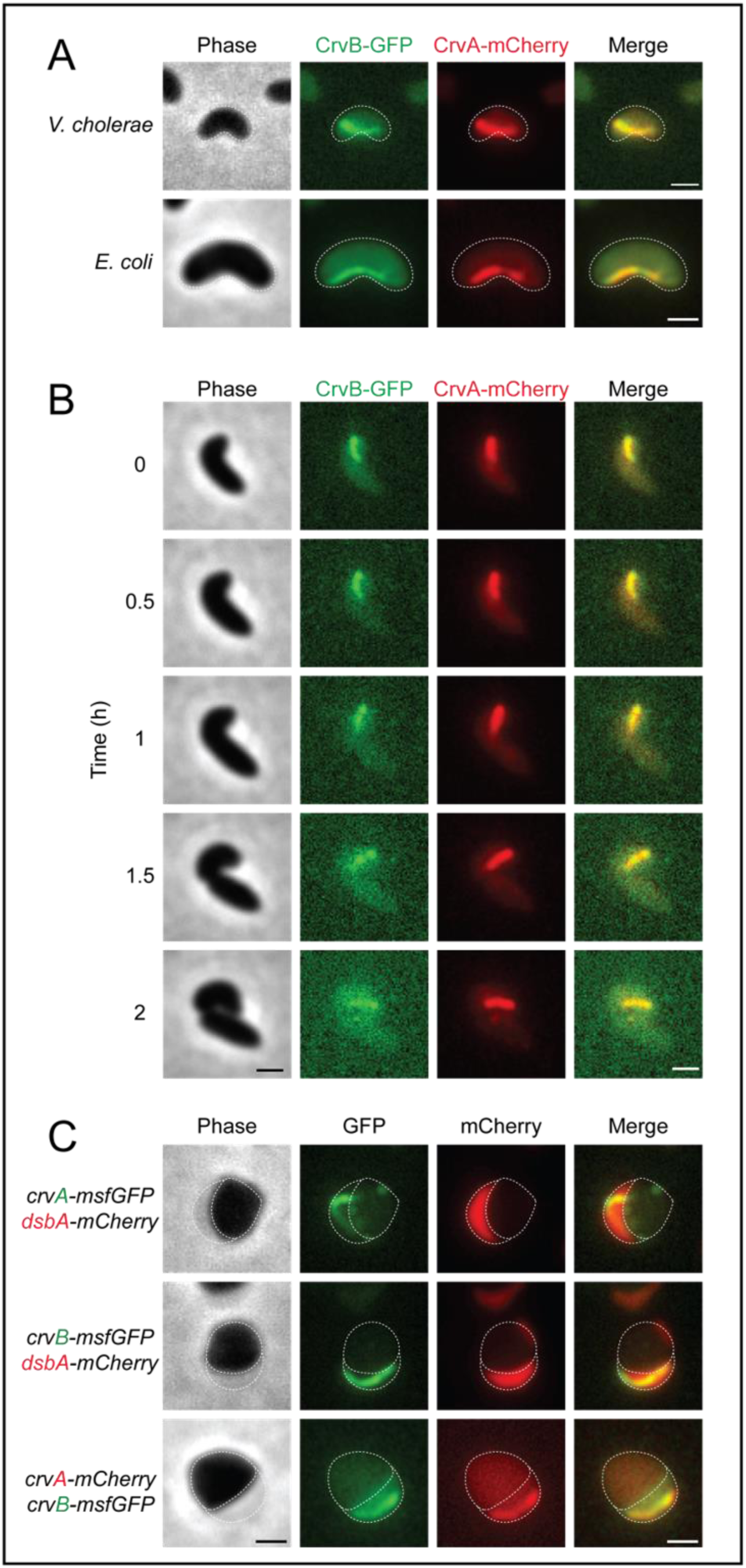
CrvA and CrvB colocalize in a periplasmic filament. (A) Curved *V. cholerae* and *E. coli* cells expressing CrvB-GFP and CrvA-mCherry fusions. (B) Time lapse images of CrvB-GFP and CrvA-mCherry structures in growing *V. cholerae*. See Movie S1 for full time lapse. (C) Fluorescent CrvA and CrvB filaments with periplasmic DsbA-mCherry in *V. cholerae* after moenomycin treatment. (A,C) Dotted lines represent outline of cell body and periplasm. Scale bars represent 1μm.

In Gram-negative bacteria, the cell wall is located between the inner and outer cell membranes in a subcellular compartment called the periplasm. In contrast to cytoskeletal elements, which assemble in the cytoplasm, CrvA is a periskeletal element that assembles in the periplasm ^14^. To determine whether CrvB is also periplasmic, we labelled the periplasm by fusing mCherry to the secretion signal of the periplasmic protein DsbA ^21^. When we perturbed these cells with the antibiotic moenomycin, cells rounded up and the periplasm visibly protruded from the cytoplasm ^22^. As expected for a periplasmic protein, CrvA-GFP filaments localized to the DsbA-mCherry-containing periplasm ^14^ (Fig. 3C). CrvB-GFP filaments also colocalized with DsbA-mCherry, indicating that CrvB is periplasmic. Furthermore, CrvA-mCherry and CrvB-GFP filaments remained colocalized in rounded moenomycin-treated cells, indicating that their colocalization to this periskeletal structure does not depend on the wild-type cellular geometry (Fig. 3C).

Next, we wondered whether CrvA and CrvB have distinct roles in assembling the periplasmic filament. Deleting *crvA* resulted in diffuse CrvB-GFP localization, indicating that CrvA is required for CrvB assembly (Fig. 4A). Deleting *crvB* also significantly disrupted CrvA-GFP, but in a qualitatively different manner. While most Δ*crvB* cells still exhibited well-defined CrvA-GFP structures, these cells typically exhibited multiple small CrvA-GFP puncta and only ∼6% had long CrvA-msfGFP filaments similar to those seen in wild type (Fig. 4A). We thus hypothesized that CrvA can form small structures on its own and that CrvB promotes the higher-order assembly of CrvA. To test this model, we placed *crvB* under arabinose-inducible control (P_bad_) at an ectopic locus and observed the effect of *crvB* expression on CrvA-GFP assembly. Consistent with our hypothesis, *crvB* expression increased CrvA-GFP assembly (Figs. 4B and 4C) as well as cell curvature (Figs. 4B and 4D) in a dose-dependent manner. Higher levels of *crvB* expression also caused earlier onset of CrvA-GFP assembly (Fig. 4C), raising the question of whether the dynamics of CrvA-GFP assembly are important for the dynamics of *V. cholerae* curvature.

**Fig. 4.**
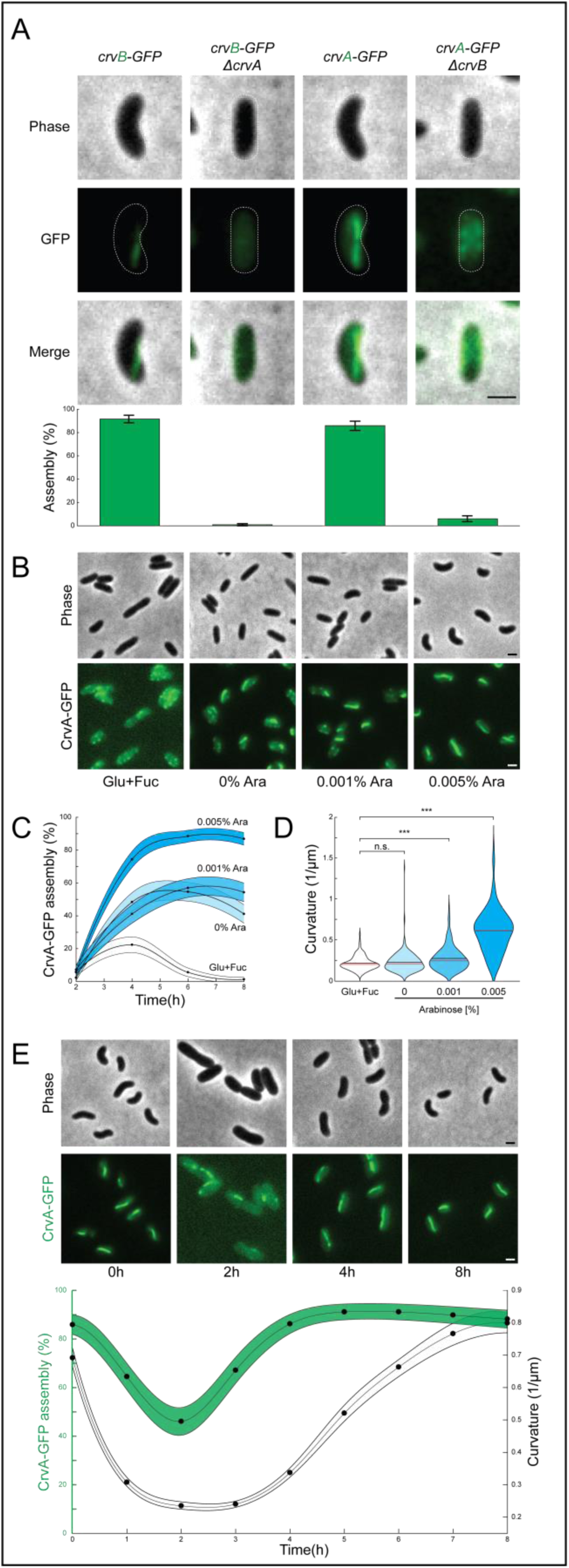
CrvA and CrvB synergize to promote filament assembly in *V. cholerae*. (A) Representative cells (top) and quantified CrvA-GFP and CrvB-GFP filament assembly (bottom, error bars indicate ± 95% confidence intervals) in deletion mutants. (B) Representative fields after repression (Glu+Fuc) and induction (Ara) of *P*_*bad*_*-crvB*. (C) Dynamics of CrvA-GFP assembly from (B) ± 95% confidence intervals. (D) End point curvature measurements from (B). (E) Dynamics of mean curvature and CrvA-GFP filament assembly ± 95% confidence intervals. (A-E) Filament assembly is the percentage of cells with fluorescent filaments similar to those in the wild type background. n=300; Wilcoxon rank-sum test; n.s.= p>0.4, ***=p<0.0001. 95% confidence intervals on filament assembly and curvature determined by Agresti-Coull method and bootstrapping, respectively. Scale bars represent 1μm.

To assess the relationship between CrvA-GFP assembly and cell curvature dynamics, we simultaneously observed CrvA-GFP assembly and cell curvature in a wild-type background. Cell curvature decreased as CrvA-GFP filaments were lost (Fig. 4E), and in these straight cells, CrvA-GFP formed multiple small structures resembling those in Δ*crvB*. Our time course also showed that cell curvature only began to increase after the formation of long CrvA-GFP filaments (Fig. 4E). Taken together these results demonstrate that the CrvAB module forms a dynamic periskeletal structure, that CrvB promotes the formation of this higher-order structure, and that the timing of its assembly correlates to the dynamics of curvature.

## Discussion

Together, our results demonstrate that both CrvA and CrvB have specialized roles in the assembly of a periskeletal structure that can impart cell shape complexity onto bacterial species separated by 2.5 billion years of evolution ^23^. The ability of CrvAB to modulate cell shape independently of species-specific growth mechanisms could have bioengineering applications like improving the stability of synthetic probiotics, as cell curvature can enhance intestinal colonization ^14^. The periskeletal nature of CrvAB allows this module to generate asymmetry similarly to cytoskeletal elements, but without requiring transmembrane interaction partners to influence periplasmic cell wall synthesis. Most cytoskeletal elements require ATP or GTP for dynamics. These energy sources are absent in the periplasm, but CrvAB demonstrate that periplasmic polymers can be dynamic nevertheless. As PG is chemically similar among bacteria ^24^, a direct interaction with the cell wall may provide an explanation for the unprecedented range of species in which CrvAB are functional. Alternatively, CrvAB may produce asymmetry by directing the activity of PG enzymes that function outside of the core cell shape complexes. In a variety of species, determinants of complex shape have been identified ^3,25^, but it remains to be seen if these systems depend on core shape machinery or if they will join CrvAB as autonomous cell shape modules. Furthermore, there are complex bacterial morphologies without characterized determinants. Perhaps these diverse forms are built not by co-option of core cell biology, but by simple cell shape modules like the one that makes *V. cholerae* curved.

## Acknowledgments

We thank members of the Gitai and Shaevitz labs for helpful discussions, the Bina and Bassler labs for reagents, Joe Sanfilippo, Matthias Koch, Courtney Ellison, Josh Shaevitz, and Tom Silhavy for helpful feedback on the manuscript.

## Author contributions

All experiments were performed by N.R.M., E.B., and T.M.B. with design input from Z.G. Data and sequence analysis were performed by N.R.M. and B.P.B. The manuscript was written by Z.G. and N.R.M. with input from the other authors.

## Data and materials availability

All data is available in the main text or the supplementary materials.

## Supplementary Materials

Figures S1-S5

Tables S1-S2

Movies S1-S2

References (27-47)

## Materials and Methods

### Bacterial strains and growth conditions

Bacterial strains and plasmids used are listed in Table S1. *Vibrio cholerae, Escherichia coli, Pseudomonas aeruginosa* were cultured in LB medium (10g/L NaCl (Sigma), 10g/L tryptone (Bacto), 5g/L yeast extract (Bacto)) at 37°C with aeration. *Agrobacterium tumefaciens* was cultured in LB medium at 30°C with aeration. *Aliivibrio fischeri* was cultured either on LB agar (Bacto) plates or in LM liquid medium (20g/L NaCl, 10g/L tryptone, 5g/L yeast extract) at 30°C with aeration. *Caulobacter crescentus* was grown in PYE medium (2g/L peptone (Bacto), 1g/L Yeast extract, 1mM MgSO_4_ (Sigma), 0.5mM CaCl_2_(Sigma)) at 30°C with aeration. Cultures in Fig 3 and Fig 4B were grown in 2mL culture volumes, while all other experiments were conducted in 5mL culture volumes. Chromosomal mutations in *V. cholerae* and *A. fischeri* were made using suicide vectors pKAS32(Skorupski and Taylor, 1996) and pRE112(Edwards et al., 1998), respectively. Plasmids were introduced by conjugal transfer from *E. coli* S17 into *V. cholerae, C. crescentus, A. tumefaciens*, and *A. fischeri.* Plasmids were introduced by electroporation into *P. aeruginosa* and *E. coli MG1655.* Antibiotics were used in the following concentrations: Kanamycin 50μg/mL (*V. cholerae, E. coli*), Carbenicillin 25μg/mL (*A.* tumefaciens), Carbenicillin 200μg/mL (*P.aeruginosa*), Chloramphenicol 1μg/mL (*C.* crescentus).

### Microscopy

Images were obtained using a Hamamatsu ORCA-Flash4.0 Digital CMOS camera on either a Nikon 90i microscope with a Nikon 100x Plan Apo NA=1.40 oil immersion objective lens or a Nikon Ti-E microscope with a Nikon 100x Plan Apo NA=1.45 oil immersion objective lens. For fluorescence, GFP-L and Cy5 HYQ filter cubes (90i) or GFP-HQ and mCherry filter cubes (Ti-E) were used to visualize msfGFP and mCherry fluorescent fusions, respectively. Excelitas X-Cite 120 LED*Boost* was used for illumination on both microscopes. All hardware was controlled by NIS Elements (version 4.60.00). Suspensions of *A. fischeri* cells were spotted on 1-2% (weight/volume) agarose(Invitrogen) pads of LM for imaging. For all other species, 1-2% agarose pads of M9 Salts (47.8mM Na_2_HPO_4_(Sigma), 22mM KH_2_PO_4_(Sigma), 8.6mM NaCl, 18.6mM NH_4_Cl(Sigma)) were used. For growth on pads, exponentially growing *V. cholerae* were spotted on 1-2% agarose pads of M9 supplemented with 0.5% glucose (Sigma) and incubated at 25°C throughout the experiment. All geometric measurements were performed on phase-contrast images at 100X magnification. For merged images, problems with filter alignment were manually corrected by adjusting contrast such that the cell body but not features within the cell were clearly visible. The image plane was then shifted so that outline of this fluorescence was aligned to the phase-contrast image or other fluorescent channel.

### Measurement of cellular geometry

Phase images were analyzed using the MATLAB script Morphometrics(Ursell et al., 2014) to segment cell contours and fit them with centerlines. Contours touching adjacent cells or having visible cleavage furrows were excluded from analysis to avoid overestimation of curvature due to the orientation of neighboring cells or recently divided cells. Curvature is root mean squared curvature of the centerline in units of inverse microns (μm^-1^). Width is mean width of the contour along the entire centerline in microns (μm). Population measurements are from pooled measurements of 100 individual cells from three biological replicates performed on different days (300 cells per measurement). To account for differences in the number of cell contours in each of the three replicates, 100 cells were randomly selected from each replicate before pooling. Violin plots represent one such randomized population. Populations that were also measured for CrvA/CrvB-msfGFP filament assembly were not randomized because only the first 100 valid contours from each replicate were analyzed; therefore, all three replicates had an equal number of contours. For statistical tests of populations, the Wilcoxon two-sided rank sum test was used. For tests on randomized populations, the random sampling and statistical testing was repeated one thousand times and the mean p-value was reported to minimize the effect of any one sub-sample. Representative cells shown in Figs 1A,C,D, Fig 4D, Figs S1-S5 are representative of the 3^rd^ quartile of length in their respective populations to facilitate visual comparison of cells with similar lengths. Unless otherwise indicated, these cells represent the 95^th^ quantile of curvature in their respective populations.

### Measurement of CrvA/CrvB-msfGFP filament assembly

Phase/GFP image stacks were split into their constituent channels and the phase channel was used to obtain cell contours using Morphometrics. After excluding touching and dividing cells as described above, the first 100 contours from each sample were superimposed on the corresponding GFP image. The GFP signal of each contour was manually inspected and categorized as “assembled” if most of the fluorescence signal formed a single, linear structure at least half the length of the cell. This was repeated on contours from three biological replicates performed on different days and pooled for population measurements (300 cells total). The Agresti-Coull method was used to estimate the 95% confidence intervals of this measurement and the center of this interval is reported as the proportion of the population that has assembled GFP structures. This proportion is referred to as the “assembly” of the population and is expressed as a percentage.

### Complementation and heterologous expression of crvA, crvB, and crvY

For *crvA* or *crvB* expression, plasmid-borne alleles were placed under the control of a tetracycline-inducible promoter(Bina et al., 2014) and the following ribosome-binding site: AGGAGCTAAGGAAGCTAAA. When expressed together, both open reading frames are under control of the same promoter separated by the same ribosome-binding site. The same expression fragment was cloned into different plasmid backbones when required for heterologous expression. Anhydrotetracycline was added at the beginning of subculture in the following concentrations with a final culture volume of 5mL: 0ng/mL for *V. cholerae, E. coli*, and *P. aeruginosa*; 2ng/mL for *A. tumefaciens*; 5ng/mL for *C. crescentus*. For *V. cholerae, E. coli*, and *P. aeruginosa* saturated overnight cultures were diluted 1:1000 into LB with appropriate antibiotics and grown for 6h before imaging. For *C. crescentus* saturated overnight cultures were diluted 1:100 into PYE with appropriate antibiotics and grown for 7h before imaging. For *A. tumefaciens* saturated overnight cultures were diluted 1:20 into LB with appropriate antibiotics and grown for 6h before imaging. For *crvY* expression in *V. cholerae* Δ*crvAB*, a plasmid-borne *crvY* allele from *A. fischeri* was placed under the control of an arabinose-inducible promoter(Guzman et al., 1995) and the ribosome-binding site above. 0.1% L-Arabinose (Sigma) was added to LB with Kanamycin at the beginning of subculture with a final culture volume of 5mL. Saturated overnight cultures were diluted to an optical density (OD_600_) of 0.001 and grown for 4h before imaging.

### Sublethal treatment with A22, Cephalexin, and Moenomycin

For moenomycin treatment saturated overnight LB cultures were diluted 1:1000 into fresh LB supplemented with 500ng/mL moenomycin and incubated at 37°C for 5-6h before imaging. Representative cells were chosen such that their orientation relative to the imaging plane allowed visual resolution of the periplasmic and cytoplasmic spaces. For A22 and cephalexin treatment, saturated overnight LB cultures were diluted 1:1000 into fresh LB and grown for 4h after which pre-treatment samples were imaged. At 4h A22 and cephalexin were added in indicated concentrations. For A22 treatment cells were grown for an additional 4h before imaging. For Cephalexin treatment, and simultaneous A22+cephalexin treatment cells were imaged 2h after drug addition.

### Titration of crvB expression in V. cholerae

*crvB* was deleted in a strain background expressing *crvA-msfGFP* from the native locus producing NM149. Into this background, a wild-type *crvB* was introduced at an ectopic locus on the opposite chromosome (*VC1378*) under the control of an arabinose-inducible promoter (Guzman et al., 1995) producing NM565. Saturated vernight cultures of NM565 were grown under repressing conditions (0.05% D-Fucose(Acros), 0.5% D-Glucose) in LB and back-diluted to an OD_600_ of 0.001 into repressing conditions or into LB supplemented with indicated concentrations of L-Arabinose. Images were taken starting 2h after back-dilution.

### Phylogenetic analysis

A multiple alignment of the unique sequences of crv proteins (all found within the Vibrionales clade) and an outgroup of the closest homologs found after excluding the Vibrionales was made using MAFFT(Katoh and Standley, 2013) with blosum80(Henikoff and Henikoff, 1992) as the scoring matrix. Following this multiple alignment, a phylogenetic tree was constructed using BMGE/FastTree(Criscuolo and Gribaldo, 2010; Guindon et al., 2010). To compare the local sequence similarity between the CrvA, CrvB, and CrvY, we divided the sequences into four clades (c ={outgroup, crvA, crvB, crvY}) and calculated a local, windowed alignment score(Capra and Singh, 2007) compared to the CrvY clade using the blosum80 scoring matrix.

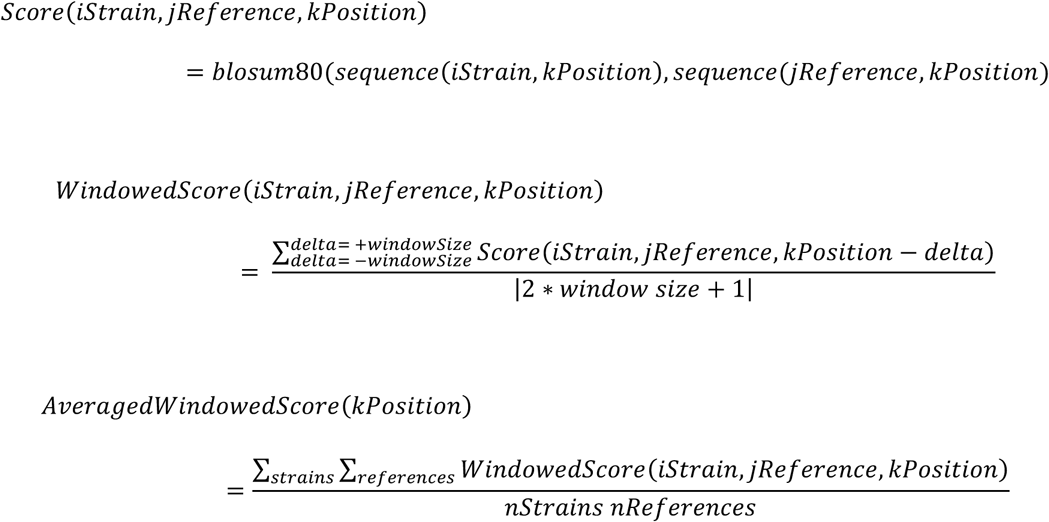

