## Supplementary Materials for "The evolution of bacterial shape complexity by a curvature-inducing module"

**Supplementary Materials include:**

Figs. S1 to S5

Table S1

Caption for Table S2

Captions for Movies S1 to S2

**Other Supplementary Materials for this manuscript include the following:**

Table S2

Movies S1 to S2

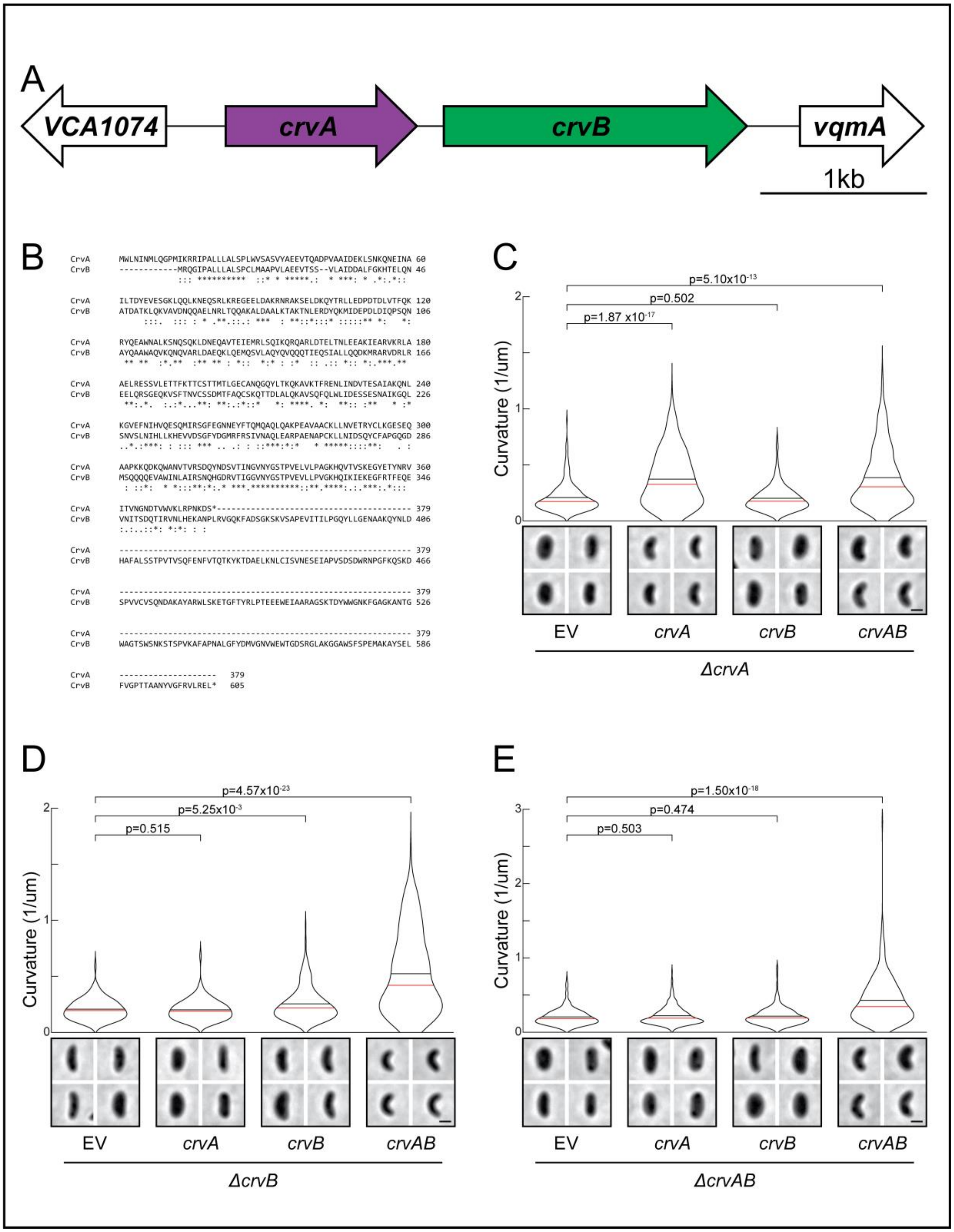


**Fig. S1 CrvAB sequence alignment and complementation analysis.** (A) The *crvA* and *crvB* open reading frames drawn to scale at the native locus in *V. cholerae*. (B) Alignment of *V. cholerae* CrvB amino acid sequence to *V. cholerae* CrvA by Clustalω[36]. (C-E) Curvature of (C)

*ΔcrvA*, (D) *ΔcrvB*, or (E) *ΔcrvAB* populations expressing an empty vector (EV) or plasmids with the indicated gene(s). (C-E) Images represent 95$^{th}$ percentile of curvature in respective populations. Scale bars represent 1μm. p-values determined by Wilcoxon rank sum test; n=300.

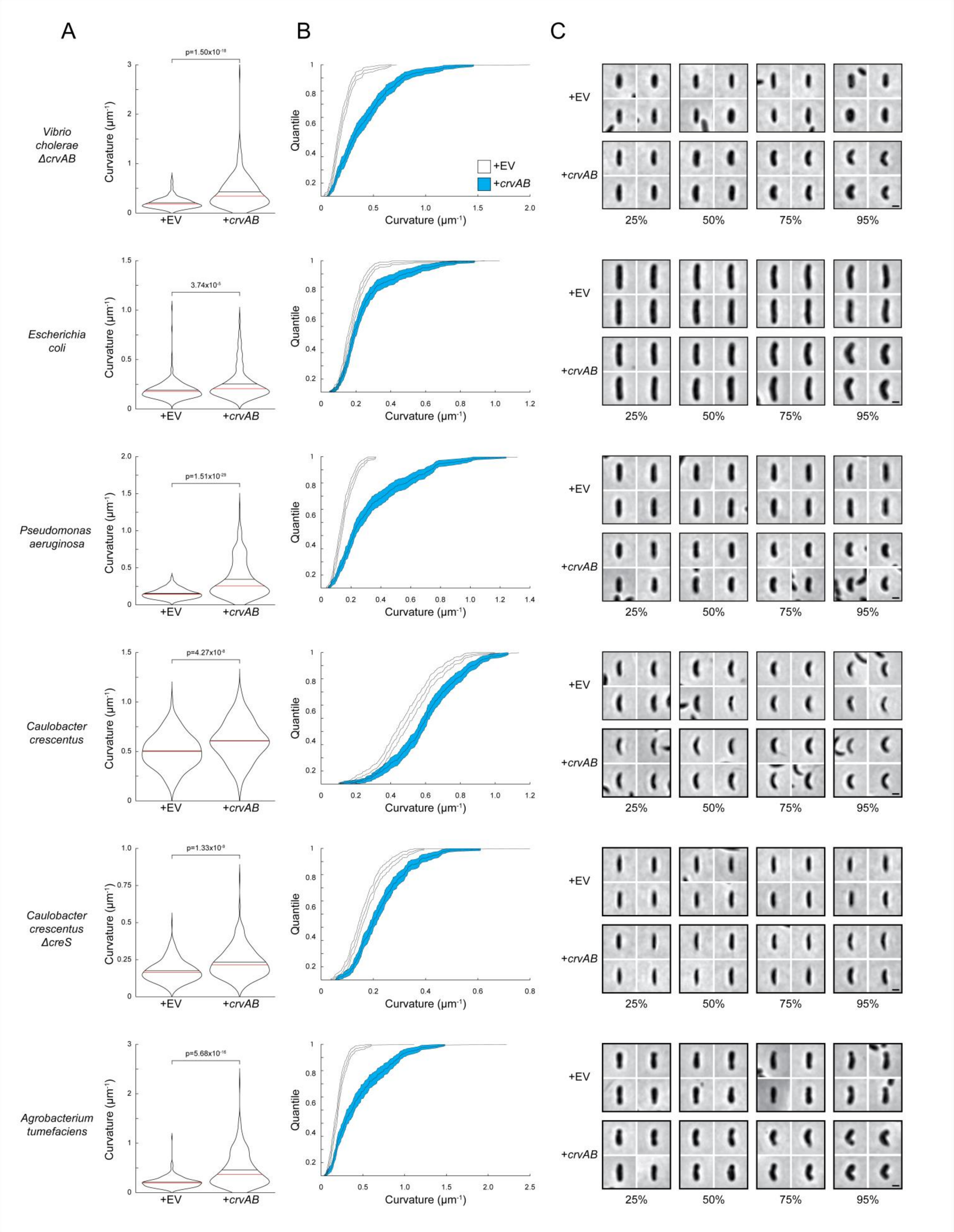

A
B
C
Vibrio cholerae ΔcrvAB
Escherichia coli
Pseudomonas aeruginosa
Caulobacter crescentus
Caulobacter crescentus ΔcreS
Agrobacterium tumefaciens
p=1.50x10-18
3.74x10-5
p=1.51x10-29
p=4.27x10-8
p=1.33x10-9
p=5.68x10-16
Curvature (µm-1)
Quantile
+EV
+crvAB
25%
50%
75%
95%

**Fig. S2 Quantification of heterologous CrvAB expression.** (A) Curvature of populations expressing an empty vector (+EV) or a plasmid with *crvA* and *crvB* (+*crvAB*). p-values determined by Wilcoxon rank sum test; n=300. (B) Cumulative distribution functions of populations in (A) ± 95% confidence intervals. (C) Images represent indicated quantiles from populations in (A). Scale bars represent 1μm. Each row corresponds to the species labeled to the left of (A).

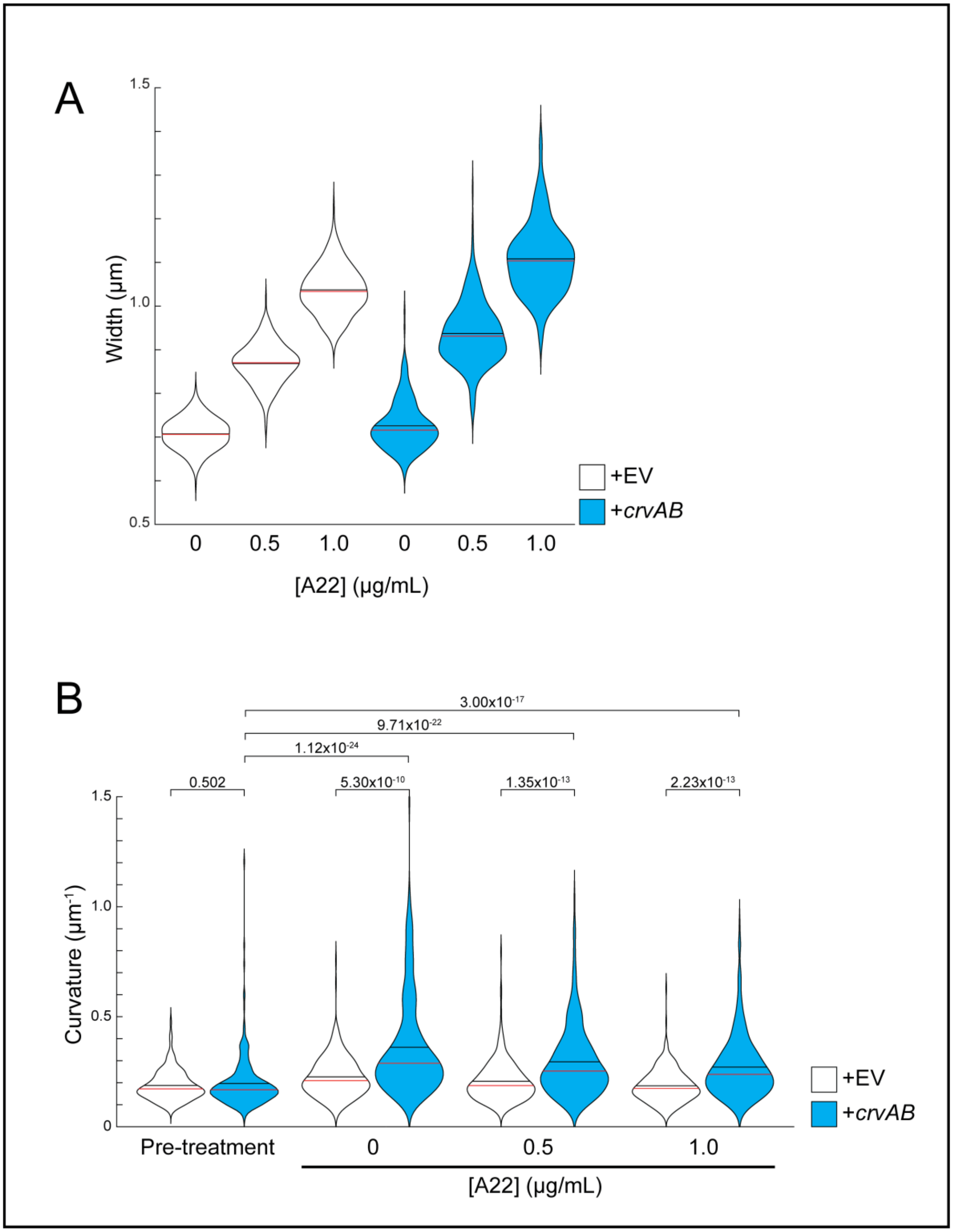


**Fig. S3 Quantification of A22 treatments.** (A) Cell width of *E. coil* populations expressing an empty vector (+EV) or a plasmid with *crvA* and *crvB* (+*crvAB*) after A22 treatment. (B)

Curvature of populations in (A) before and after A22 treatment. p-values determined by Wilcoxon rank sum test; n=300.

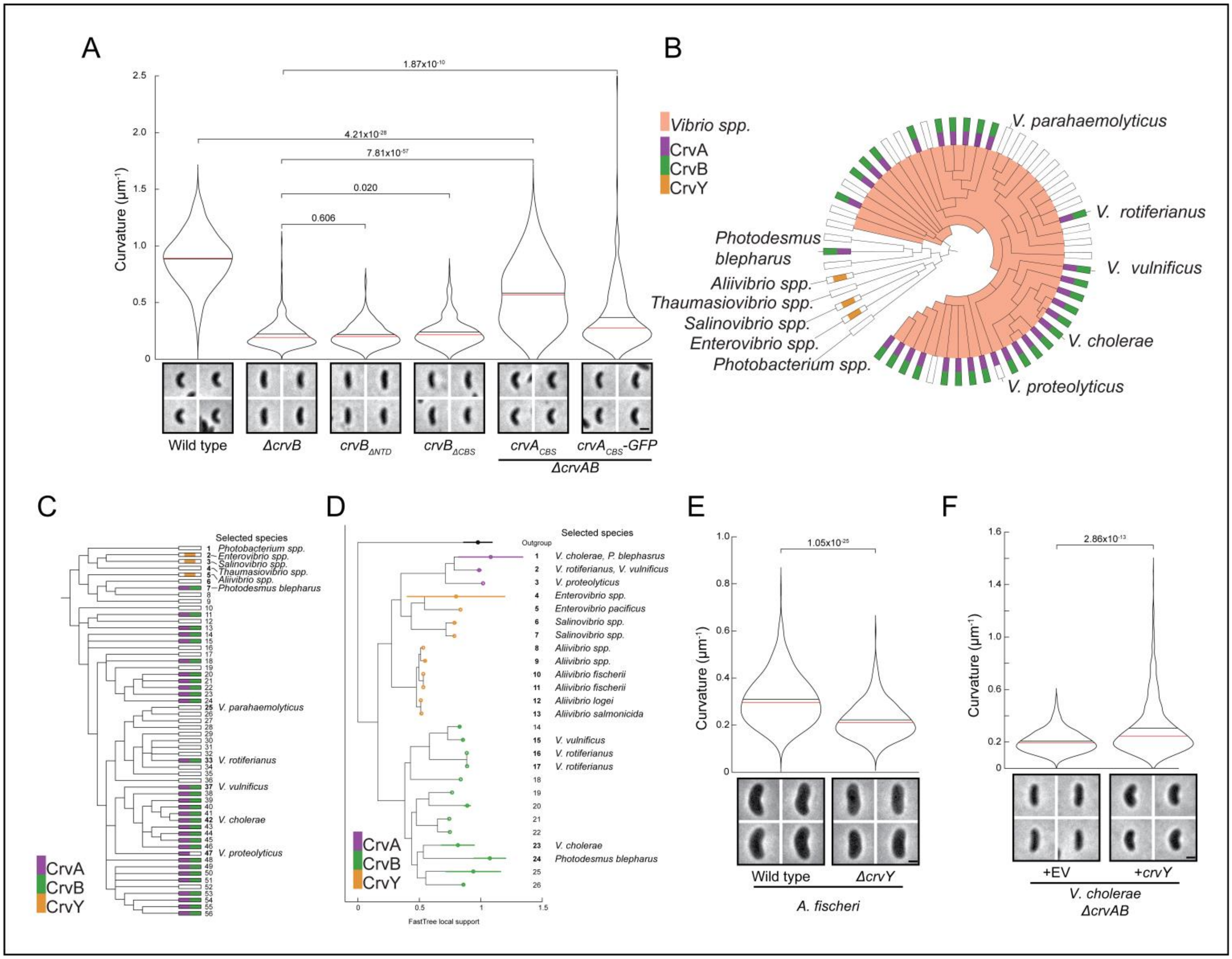


**Fig. S4 Structure-function analysis and phylogeny of hybrid Crv proteins.** (A) Curvature of populations of *V. cholerae* expressing the indicated *crvB* truncations or *crvA*$_{CBS}$ chimeras from the native *crvB* and *crvA* loci, respectively. Images taken from saturated overnight cultures. (B) Extended cladogram from Fig 1B including non-vibrio species and CrvY homologs. Selected species are indicated. (C) Linear form of (B). (D) Phylogeny of all sequenced CrvA, CrvB, and CrvY homologs. Terminal nodes are placed at the mean ± standard deviation of the sequences collapsed into each node. (E) Curvature of populations of *A. fischeri* wild type or *ΔcrvY* after colonies were suspended and grown in LM for 4h at 30°C. (F) Curvature of populations of *V.*

*cholerae ΔcrvAB* expressing an empty vector (+EV), a plasmid with *crvY* from *A. fischeri*. (A,E,F) Images represent $95^{th}$ percentile of curvature in respective populations. Scale bars represent 1µm. p-values determined by Wilcoxon rank sum test; n=300. (C,D) Numbers are “Clade IDs” for reference to Table S2, which contains full composition of terminal nodes.

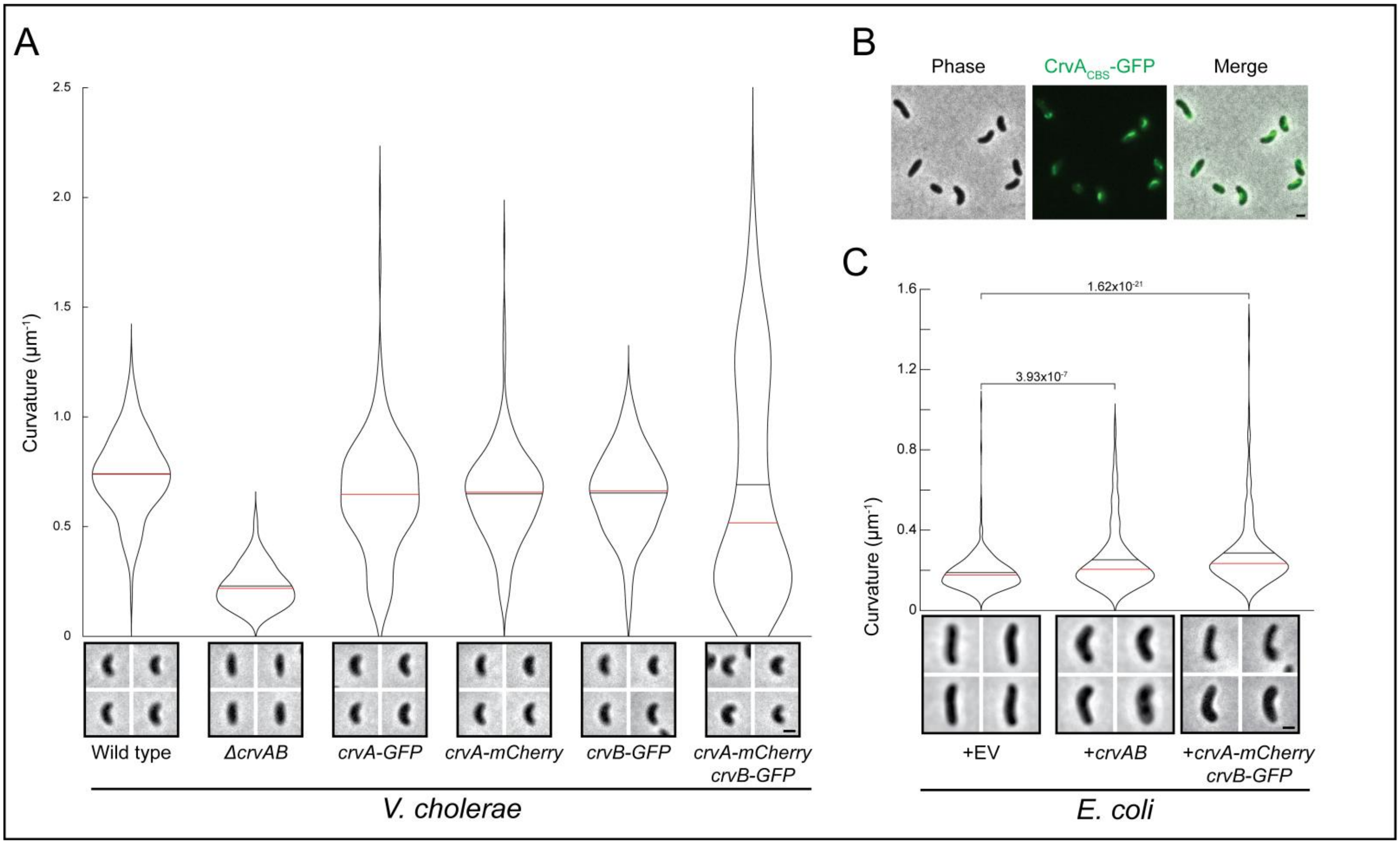


**Fig. S5 Functionality of CrvAB fluorescent fusion proteins.** (A) Curvature of populations of *V. cholerae* expressing the indicated fusion proteins from the native genomic locus. Data from wild type and *ΔcrvAB* are repeated from Fig 1A. (B) Representative image of cells expressing the *crvA*$_{CBS}$ chimera tagged with msfGFP (CrvA$_{CBS}$-GFP). (C) Curvature of *E.coli* populations expressing an empty vector (+EV), a plasmid with *crvA* and *crvB* (+*crvAB*), or the same plasmid with *crvA-mCherry* and *crvB-GFP*. Data from +EV and +*crvAB* are repeated from Fig S3. (A,C) Images were taken 6h after 1:1000 dilution of saturated overnight cultures. Images represent 95$^{th}$ percentile of curvature in respective populations. p-values determined by Wilcoxon rank sum test; n=300 (A-C) Scale bars represent 1µm.

**Table S1.** Bacterial strains and plasmids used in this study.

| Strain | Description | Source |
|---|---|---|
| ZG1307 | (*Vc*) C6706 | [37] |
| S17-1 | *E. coli* used for cloning and plasmid conjugal transfer | [38] |
| NM005 | (*Vc*) C6706 Δ*crvA* | [14] |
| NM146 | (*Vc*) C6706 Δ*crvB* | This study |
| NM189 | (*Vc*) C6706 Δ*crvAB* | This study |
| NM443 | (*Vc*)Δ*crvA*+pEVS143::$P_{tet}$-*crvA* | This study |
| NM444 | (*Vc*)Δ*crvB*+pEVS143::$P_{tet}$-*crvA* | This study |
| NM445 | (*Vc*)Δ*crvAB*+pEVS143::$P_{tet}$-*crvA* | This study |
| NM288 | (*Vc*)Δ*crvA*+pEVS143 EV | This study |
| NM335 | (*Vc*)Δ*crvB*+pEVS143 EV | This study |
| NM290 | (*Vc*)Δ*crvAB*+pEVS143 EV | This study |
| NM607 | (*Vc*)Δ*crvA*+pEVS143::$P_{tet}$-*crvAB* | This study |
| NM608 | (*Vc*)Δ*crvB*+pEVS143::$P_{tet}$-*crvAB* | This study |
| NM510 | (*Vc*)Δ*crvAB*+pEVS143::$P_{tet}$-*crvAB* | This study |
| ZG799 | (*Ec*) MG1655 | [39] |
| NM609 | (*Ec*)MG1655+pEVS143::$P_{tet}$-*crvAB* | This study |
| NM617 | (*Ec*) MG1655+pEVS143 EV | This study |
| NM672 | (*Pa*)PA14 | [40] |
| NM673 | (*Pa*)PA14+pUCP18::$P_{tet}$-*crvAB* | This study |
| NM674 | (*Pa*)PA14+ pUCP18 EV | This study |
| ZG66 | (*Cc*)CB15N | [41] |
| NM803 | (*Cc*)CB15N+pRXMCS-6 EV | This study |
| NM804 | (*Cc*) CB15N+pRXMCS-6::$P_{tet}$-*crvAB* | This study |
| ZG1539 | (*Cc*)CB15N Δ*creS* | [42] |
| NM692 | (*Cc*)Δ*creS*+pRXMCS-6 EV | This study |
| NM693 | (*Cc*) Δ*creS*+pRXMCS-6::$P_{tet}$-*crvAB* | This study |
| NM697 | (*At*) C58 | [43] |
| NM701 | (*At*)C58+pRXMCS-6 EV | This study |
| NM702 | (*At*)C58+pRXMCS-6::$P_{tet}$-*crvAB* | This study |
| NM583 | (*Vc*)*crvA-msfGFP* | This study |
| NM162 | (*Vc*)*crvA-mCherry* | This study |
| NM147 | (*Vc*)*crvB-msfGFP* | This study |
| NM169 | (*Vc*) *crvA-mCherry*;*crvB-msfGFP* | This study |
| NM682 | (*Ec*)MG1655+pEVS143::$P_{tet}$-*crvA-mCherry-crvB-msfGFP* | This study |
| NM613 | (*Vc*)*crvA-msfGFP*+pEVS143::$P_{tet}$-$SS_{VcdsbA}$-*mCherry* | This study |
| NM614 | (*Vc*)*crvB-msfGFP*+pEVS143::$P_{tet}$-$SS_{VcdsbA}$-*mCherry* | This study |
| NM148 | (*Vc*) *crvB-msfGFP*; Δ*crvA* | This study |
| NM149 | (*Vc*) *crvA-msfGFP*; Δ*crvB* | This study |
| NM565 | (*Vc*) *crvA-msfGFP*; Δ*crvB*; *VC1378*::$P_{bad}$-*crvB* | This study |

| | | |
|---|---|---|
| NM859 | (*Vc*) *crvBΔNTD* (Δ24-359) | This study |
| NM853 | (*Vc*) *crvBΔCBS*(Δ360-605) | This study |
| NM353 | (*Vc*) Δ*crvB*; *crvA-CBS*(*crvB* 360-605) | This study |
| NM584 | (*Vc*) *crvA-CBS-msfGFP* | This study |
| NM329 | *A. fischeri* ES114 | [44] |
| NM814 | *A. fischeri ΔcrvY* | This study |
| NM527 | Δ*crvAB*+pEVS143::$P_{bad}$-*crvY* | This study |
| Plasmid | Description | Source |
| pKAS32 | Allelic exchange vector used for *V. cholerae* chromosomal mutations | [27] |
| pRE112 | Allelic exchange vector used for *A. fischeri* chromosomal mutations | [28] |
| pEVS143 | Expression vector used in *V. cholerae* and *E. coli* | [45] |
| pUCP18 | Expression vector used in *P. aeruginosa* | [46] |
| pRXMCS-6 | Expression vector used in *C. crescentus* and *A. tumefaciens* | [47] |

(*Vc*): *V. cholerae*, (*Ec*): *E. coli*, (*Pa*): *P. aeruginosa*, (*Cc*): *C. crescentus* (*At*): *A. tumefaciens*, EV: Empty vector

**Table S2(separate file).** Genomic accession numbers, crv protein accession numbers, and clade labels for each of 921 Vibrionacae genomes. Clade labels are listed in figures S4C and S4D. Excel file with two header rows and 921 data rows. Column names (first header row) and descriptions of the data in each column (second header row) are included for each of the 17 columns.

**Movie S1.**

Time lapse of intact CrvA-mCherry/CrvB-GFP structure in *V. cholerae* cell from Fig 3B. The fluorescent structure remains intact throughout the time course and remains associated with the curved daughter cell from the initial division. White arrowhead indicates the position of a fluorescent filament. Images were captured at a rate of 1frame/5min. Scale bar is 2μm.

**Movie S2.**

Time lapse of intact CrvA-mCherry/CrvB-GFP structure in hyper-elongated *V. cholerae.* The indicated structure is in a cell that is spontaneously filamenting by growing without cell division. A single, long fluorescent structure can be seen travelling along with the cell and rotating around its surface as the cell grows across the field of view. White arrowhead indicates the position of a fluorescent filament. Images were captured at a rate of 1frame/5min. Scale bar is 2µm.
